# From Pan-Life Phase Insights to PhaseHub: Analyzing Condensate Complexity

**DOI:** 10.1101/2025.05.15.653974

**Authors:** Qiyu Liang, Weibo Gao, Yansong Miao

**Affiliations:** School of Physical and Mathematical Sciences, Nanyang Technological University, 637371, Singapore; School of Biological Sciences, Nanyang Technological University, 637551, Singapore; School of Electrical and Electronic Engineering, Nanyang Technological University, 639798, Singapore; Institute for Digital Molecular Analytics and Science, Nanyang Technological University, 636921, Singapore

## Abstract

Macromolecular condensation is crucial in biological signaling pathways that regulate development and adaptation processes. However, lacking conserved residues in disordered region of condensate proteins hampers alignment-based analyses of their evolutionary trajectory. To gain a comprehensive overview of phase-separating protein evolution across the Tree of Life, we analyzed phase-separating proteins of proteomes from 1,106 species. Eukaryotes and prokaryotes showed high contrast in need for phase-separating proteins and with a clear genome size-dependent correlation. Evolutionary success appears to hinge on the balance between functional condensation and avoiding harmful aggregation-prone biophysical signatures, such as certain amino acid homorepeats embedded in molecular grammar. We also identified potential phase separation-regulated signaling hubs across kingdoms by integrating phase-separation-positive proteins with protein abundance and interactome data across four model eukaryotic species. Consequently, we created PhaseHub (https://phasehub.sbs.ntu.edu.sg/), a user-friendly interface that explores key scaffold proteins and binders to pinpoint associated partners and potential multicomponent phase separation hubs.

**Teaser:** Analyse phase separation behavior of 1,106 species across kingdoms on proteome level reconstruct the evolutional history of protein condensation.

## Introduction

As a fundamental regulatory mechanism, phase separation (PS) is the key to coordinating cellular activities. Different from the rigid lock-and-key interaction model, which relies on precise structural complementarity between binding partners, the formation of PS is highly mediated by biophysical features of weak and multivalent interactions (*1, 2*), including the pi-pi/pi-cation interaction (*3*), intrinsically disordered region (IDR) (*4*), charge (*5*), hydrophobicity (*6*), etc. These flexible interactions enable biomolecular condensates to form dynamic, tunable assemblies that respond to microenvironment, expanding functional diversity beyond the constraints of classical stereospecific models (*7*). While the functional evolution of the structured domain can be dissected by multiple sequence alignment (MSA) to get residue-specific information, the evolutionary analysis of functional selection of PS systems cannot be systematically investigated using the same method. Due to the possession of highly flexible conformation with frequently occurring adaptive binding ability (*8, 9*), the amino acid sequences of IDRs are highly unconserved, restraining the usage of MSA (*10, 11*). However, although the prevalent of IDRs in phase-separated proteins (PSPs) (*12, 13*) limited its large-scale analysis, the biophysical features-based molecular grammar, such as charge distribution, hydrophobicity, and pattern combinations, are preserved within IDR and served as functional signatures throughout evolution (*14, 15*). These conserved features enable systematic analysis of functional selection in PSPs in evolution by classifying their biophysical properties among the vast set of unstructured PSPs. As a fundamental regulatory mechanism, PS is the key node to coordinating cellular processes with or without domain-based activities.

Experimental validation of PS in the living system is low-throughput and time-consuming, chemical precipitation, size exclusion, and density gradient ultracentrifugation-based methods combined with high-resolution quantitative mass spectrometry in living cells enable the discovery of *in vivo* PS protein on a larger scale (*16–18*). However, global analysis of the PSP atlas across the Tree of Life to understand the functional evolution of PS and predict heterotypic PS within specific living systems for investigating cellular functions remains a significant challenge. Addressing these challenges would provide a system-level view of biomolecular interactions and condensations, predicting cascade signaling mechanisms and their intricate crosstalk. Recently, the combination of protein interactome data with computational prediction revealed the composition of heterogeneous condensation (*19*), demonstrating the power of in silico screening on large-scale identification in the post-omics era. To understand the PS systematically across all the kingdoms of life, we applied the MolPhase prediction (*12*) on the proteome of 1106 species from six kingdoms. PSP ratio differentially appeared between prokaryotic and eukaryotic species, with the latter possess significant higher ratio of PSP. The increase of PSP is correlated with genome enlargement. Global multiple features analysis, such as amino acid homorepeats, support the formation of PS also concerted expansion with the biophysical features. We identified key functional nodes across different kingdoms by integrating quantitative proteomic and interactomic data from four model species, including *Saccharomyces cerevisiae*, *Plasmodium falciparum*, *Homo sapiens,* and *Arabidopsis thaliana*. Subsequently, we developed PhaseHub, an integrated database designed to analyze associative partners for exploring multicomponent condensates and poly-amino acid repeats, a unique PS feature that is less understood. In summary, our system-level approach integrates pan-life phase separation analysis, interactome, and expression data into a multicomponent condensation analyzer, enhancing our ability to identify and study functional signaling hubs arising from molecular condensation.

## Results

### Eukaryotes harbor extensive PS potential, contrasting prokaryotic scarcity

To gain a system view of homotypic PS and their distribution across the Tree of Life, we retrieved the whole proteome of 1106 representative species from six kingdoms of life, including 132 archaea, 364 bacteria, 159 fungi, 133 protists, 181 animals and 137 plants (Table S1). From the represent species of six kingdom (Figs. 1A-F), we found out that the MolPhase-predicted PS-negative proteins were dominated in the prokaryotic species such as the representative *Methanocaldococcus jannaschii* (archaea) and *Escherichia coli* (bacteria) (Figs. 1A-B) and reported phytopathogenic bacteria *Xanthomonas campestris* and *Pseudomonas syringae* (*12*). In contrast, the eukaryotic species showed bipolar distribution, with PS-positive protein possessing equal or even higher weightage in *Saccharomyces cerevisiae* (fungi), *Plasmodium falciparum* (protist), *Homo sapiens* (animal), and *Arabidopsis thaliana* (plant) (Figs. 1C-F). As the MolPhase score could reflect the probability of homotypic PS, we focused on the highly confident part in which MolPhase prediction scores higher than 0.9 as highly phase separation positive (HPS-Pos) and lower than 0.1 as highly phase separation negative (HPS-Neg). From the MolPhase score of the whole proteome, prokaryotic species exhibit a low HPS-Pos ratio but a high HPS-Neg ratio compared with eukaryotic organisms (Figs. 1G, H). Analysis of species within each kingdom reveals that the HPS-Pos to HPS-Neg ratio varies among kingdoms but fundamental differences between prokaryotic and eukaryotic organisms persist. For example, the HPS-Neg ratio in archaea was higher than in bacteria but not significant in the HPS-Pos ratio, and plants have a lower HPS-Pos ratio compared with fungi and protists (Figs. 1I, J). Overall, the distinct difference in the abundance of phase separation-prone proteins between prokaryotic and eukaryotic species suggests an increasing demand for more complex macromolecular interactions in the biological processes of eukaryotes throughout evolution.

**Figure 1.**
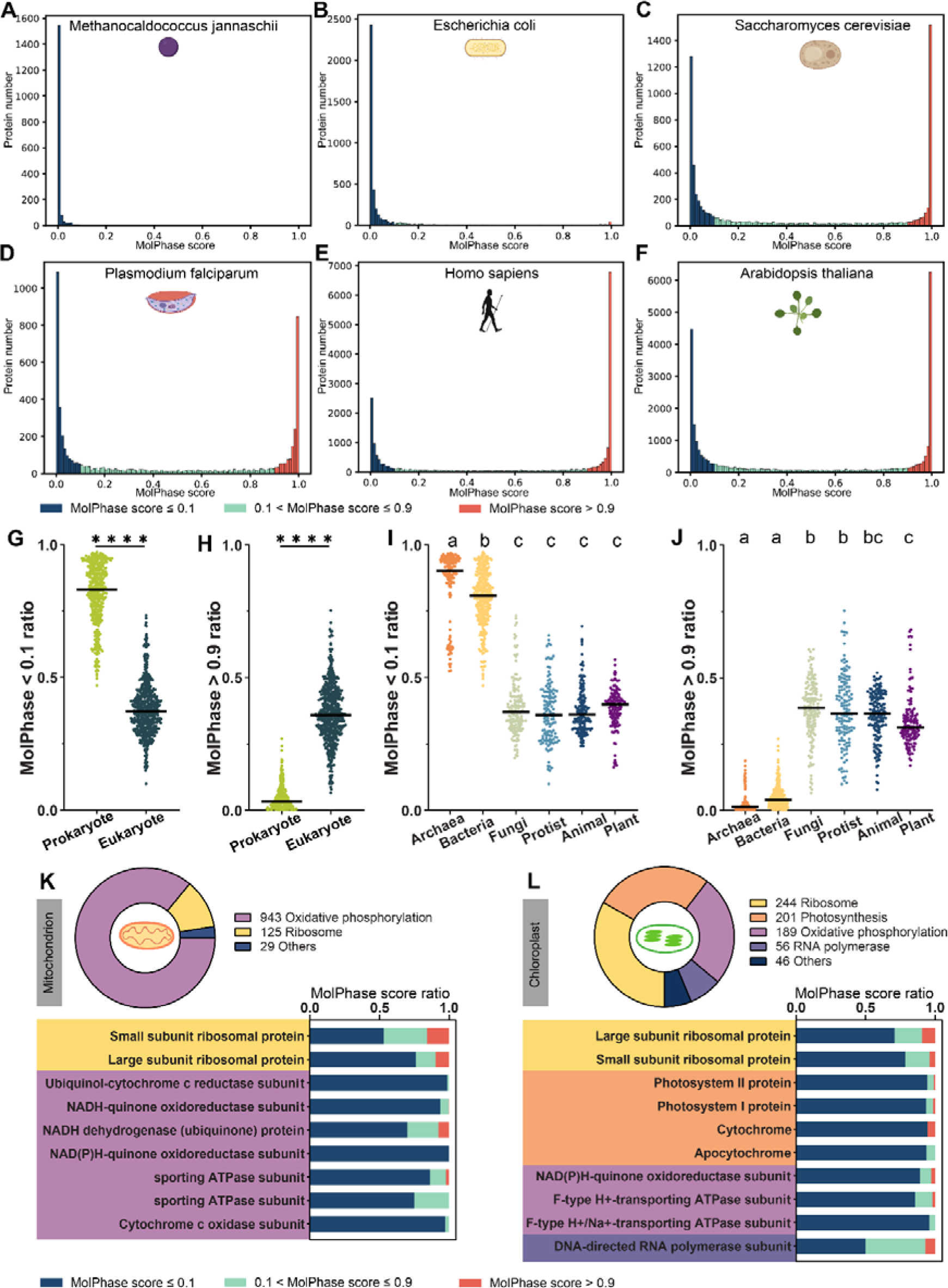
Phase separation distribution atlas of six kingdoms of life. (A)-(F) Frequency distribution histogram of MolPhase whole proteome prediction of (A) archaea *Methanocaldococcus jannaschii*, (B) bacteria *Escherichia coli*, (C) fungi *Saccharomyces cerevisiae*, (D) protist *Plasmodium falciparum*, (E) animal *Homo sapiens*, and (F) plant *Arabidopsis thaliana*. For proteins’ MolPhase prediction scores lower than 0.1 were categorized as highly phase separation negative (HPS-Neg) and prediction scores higher than 0.9 were seen as highly phase separation positive (HPS-Pos). (G)-(H) (G) HPS-Neg ratio and (H) HPS-Pos ratio comparison within the whole proteome of prokaryote species and eukaryote species. A two-tailed Mann-Whitney test carried out the statistical significance test and statistical significance is indicated as **** p ≤ 0.0001. (I)-(J) (I) HPS-Neg ratio and (I) HPS-Pos ratio comparison within the whole proteome of six kingdoms. Statistical analysis was done by one-way ANOVA with Tukey’s HSD test (p ≤ 0.05). For any two groups of data, the same letter label indicated no significant difference (p > 0.05) between the two groups. For (G)-(J), the line on top of the dots indicated the mean value. Protein pathway percentage and their relative protein family and MolPhase prediction score ratio in all mitochondrion encoded proteins. Related to Figure S1A-D. Protein pathway percentage and their relative protein family and MolPhase prediction score ratio in all chloroplast encoded proteins. Related to Figure S1C.

### Organelle-encoded proteins exhibit minimal phase separation propensity

Next, we compared the genomes of mitochondria, chloroplasts, and the nuclear genome. Mitochondria and chloroplasts retain prokaryotic genomic features, whereas the nuclear genome exhibits hallmarks of eukaryotic evolution (*20*). As mitochondrion and chloroplast evolved millions of years ago from the engulfment of aerobic and photosynthetic bacteria via endosymbiosis (Fig. S1E), the PS pattern of prokaryote were kept in mitochondrion and chloroplast even after long-term evolution in eukaryotic cell. We found that mitochondrial and chloroplastic-encoded proteins exhibited significantly higher HPS-Neg ratios than their respective nucleus-encoded protein (Figs. S1A-D). Interestingly, although another cellular plastids apicoplasts were derived from eukaryote, which evolved from secondary engulfment of plastid-contained algae (*21, 22*) (Fig. S1E), also primarily encode HPS-Neg proteins (Fig. S1D). These HPS-Neg proteins in mitochondria and plastids have maintained essential functions throughout evolution as semi-independent systems within eukaryotic cells without developing a propensity for PS. This suggests that their functional roles have remained consistent, and they have not evolved into systems prone to PS despite being part of the eukaryotic cellular environment.

Our following gene ontology biological functional analysis of mitochondrion and chloroplast-encoded proteins revealed high conservation on essential pathways, including ribosome, oxidative phosphorylation pathways, and additional photosynthesis for chloroplast (Figs. 1K, L). The mitochondrial respiratory chain supercomplex and photosystem-antenna supercomplex strongly exclude PSP to form the supermolecular complex (Figs. 1K, L), indicating that electron passage prefers tightly controlled stoichiometric interactions for function. To be noted, although chloroplast’s photosynthesis pathway has abundant HPS-Neg proteins via such homotypic phase separation analysis approach, chloroplast is also known to regulate carbon dioxide fixation via heterotypic phase separation, such as condensation of Rubisco protein complex via additional linker proteins (*23, 24*), and mitochondrial nucleoids also could self-assembly and size control via PS to modulate transcription (*25*), suggesting the necessarity of multicomponent heterotypic PS system even within a HPS-Neg dominant environment.

### PS potential correlates with genome size in small, but not large

Next, the massive expansion of PSPs from prokaryote to eukaryote motivated us to ask whether the expansion is phylogenetic distance-dependent. Then, we classified the eukaryotic species of four major kingdoms into respective clades and analyzed their HPS-Pos/Neg ratio (Fig. S2). Although we found that the most derived lineage in aminals, Chordata, possesses the most abundant HPS-Pos protein in animals and the most basal lineage in animals, Porifera, shows the lowest ratio of HPS-Pos (Fig. S2C), the propensity in phase-separation prone in eukaryotes does not always strictly correlate with phylogeny. For example, in the plant kingdom, angiosperm and gymnosperm have a higher ratio of HPS-Neg protein compared with the green algae chlorophyte (Fig. S2D), indicating that overall PS in each species does not simply follow the evolution hierarchy.

Next, we ask whether the differential PS in prokaryotes and eukaryotes could be attributed to a transition threshold, where the involvement of PS becomes sufficient to orchestrate the complexity of development and resilience, then no further PS expansion is required. We collected the genome size information of different species and analyzed their correlations with the HPS-Pos ratio. Strikingly, in prokaryotes, the enlargement of genome size shows a strong correlation with the HPS-Pos ratio, whereas in eukaryotes, this correlation is not significant (Figs. 2A, B). This finding suggests that PS is critical in providing the necessary regulatory mechanisms to maintain cellular activities when genome size expands in prokaryotes, even though the PS ratio is relatively lower in prokaryotes than in eukaryotes. And HPS-Neg ratio decreasing with the enlargement of genome size in prokaryotes and not correlated in eukaryotes (Figs. 2E, F). Furthermore, given the wide range of genome sizes in eukaryotes (from 2.3 to 40,100 Mb in our dataset), we focused on eukaryotic species with genome sizes similar to those of prokaryotes to determine the correlation between PS and genome size in this subset. Surprisingly, we found that in terms of the eukaryotes whose genome size is smaller than 20 Mb, their genome size showed a clear correlation with the HPS-Pos ratio (Fig. 2C) and even had similar fitting r^2^ compared with prokaryotes (Fig. 2A). It suggests the demand for a higher propensity for phase separation in species with the relatively greater genome when genome sizes are smaller than 20 Mb. As sufficient PSPs evolve, the percentage of HPS-Pos proteins reaches a balanced ratio, ensuring overall functionality even as genome size expands. However, even at a similar genome size, eukaryotes still process a higher ratio of HPS-Pos proteins and a lower ratio of HPS-Neg proteins (Figs. 2D, H), suggesting that the fundamental cellular structure differences allow eukaryotes to maintain more PSPs to orchestra functional life activities.

**Figure 2.**
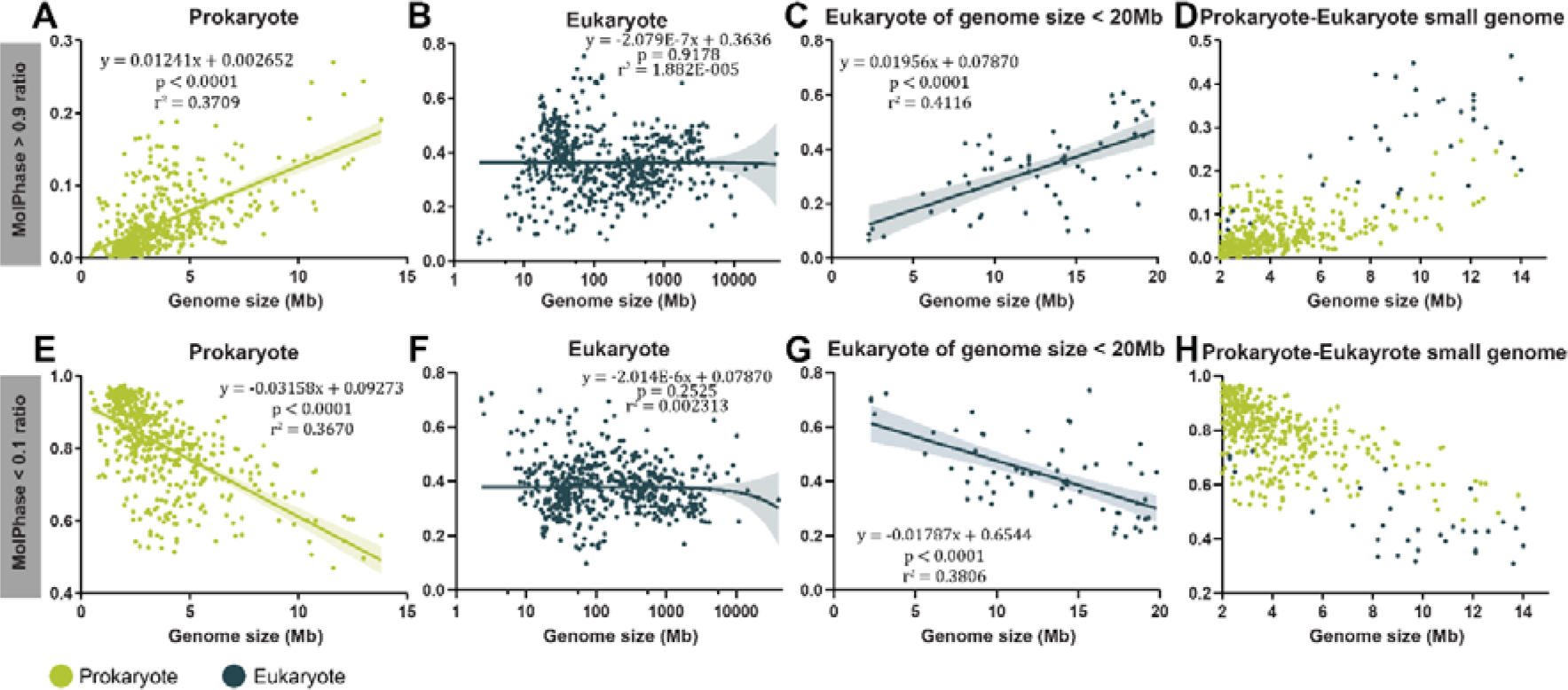
The concerted expansion and contraction of genome size and PSP ratio. (A)-(C) and (E)-(G) Linear regression fitting of genome size and HPS-Pos ratio in (A) prokaryote, (B) eukaryote, (C) eukaryote of genome size < 20 Mb. Linear regression fitting of genome size and HPS-Neg ratio in (E) prokaryote, (F) eukaryote, (G) eukaryote of genome size < 20 Mb. (D) and (H) (D) HPS-Pos protein ratio and (H) HPS-Neg protein ratio comparison of prokaryote and eukaryote species of small genome. For (A)-(C) and (E)-(G), the equation indicated the fitting results, the p-value represented the statistical significance test for slope non-zero, r^2^ indicated the coefficient of determination, the semitransparent area under the fitting line stated the 95% confidence interval of the best-fit line. For (B) and (F), the x-axis was under log10 transformation, while the fitting curve and equation were calculated based on actual data without log10 transformation. For (D) and (H), in terms of the dataset used in this study, the smallest eukaryotic genome size is 2.3 Mb, and the largest prokaryotic genome size is 13.8 Mb. Therefore, we selected species with genome sizes ranging from 2 to 14 Mb for analysis.

### Distinct biophysical features for functional PS in eukaryotes and prokaryotes

We next ask when HPS-Pos increases or reaches a balanced ratio in eukaryotic species, whether the leading biophysical signatures of PS contribute equally or differently. Pi interaction, IDR percentage, glycine ratio, prion-like domain (PLD) likelihood, sequence length, and low complexity region (LCR) percentage were analyzed, which are the top six PS promoting features according to MolPhase training set (Fig. S3A) (*12*). First, we set the feature scores of the 80^th^ percentile of *Saccharomyces cerevisiae* as standard to examine all 1106 species (Figs. S3B-G). Those feature score higher than 80^th^ percentile of *S. cerevisiae* were designated as distinct feature. And the distinct features were more prevalent in eukaryotic proteomes than in prokaryotes (Figs. S4A-F). All the analyzed feature ratios are positively correlated to the increase of the HPS-Pos ratio in the proteome (Figs. S4G-R). In addition, features concur following initial genome size expansion, such as in prokaryotic species and eukaryotes whose genome size is smaller than 20 Mb (Figs. S5A-L). Generally, the ratio of most distinct features is not correlated with genome size in overall eukaryotic species, except for PLD likelihood, which showed a weak correlation with the genome size (Figs. S5M-R). An additional analysis of average feature scores in HPS-pos proteins from prokaryotes and eukaryotes revealed distinct pattern differences. Further analysis indicated that differences exist even in terms of HPS-Pos protein between prokaryote and eukaryote. We observed stronger pi interactions in eukaryotes, longer IDRs, longer overall sequences, and a higher prevalence of PLDs (Figs. S6A-B, D-F), whereas glycine and LCRs were more prevalent in prokaryotic proteins (Fig. S6C, F). And such kind of difference is the intrinsic property of prokaryote and eukaryote, while not related to the genomel size cutoff (Figs. S6). These differences in feature usage suggest the difference of molecular gramma of PSP between prokaryote and eukaryote. Taken together, analysis of PS feature suggests different physiochemical complexity and molecular grammar for evolving functional condensation across different kingdoms.

### PSPs are more stringently regulated than non-PSPs

We further analyzed the presence and patterns of amino acid homorepeats, an LCR subclass that correlates with PS propensity (*26*). These homorepeat-containing proteins (HRPs) play critical roles in regulating molecular structures and interactions, driving evolutionary processes, or promoting higher-order assemblies (*26, 27*). Here, we defined amino acid homorepeat as containing at least five consecutive identical amino acids (*26*). HRP analysis across different species revealed that eukaryotes have a higher ratio of HRP for all 20 amino acids and longer homorepeats than prokaryotes (Fig. 3A). Among the most common homorepeats, polyD, polyS, polyN, and polyQ tend to maintain long repeat in the proteome, particularly in super-long homorepeats (>30 amino acids) (Fig. 3A). Although polyA ranks first in absolute number among both prokaryotic and eukaryotic HRP (Fig. 3B), relatively few of them form super-long homorepeats (Fig. 3A). Many HRPs in prokaryote, especially polyA, polyL containing sequences, are HPS-Neg (Fig. 3B), despite HRPs are hot candidates for PSPs. Examation of four eukaryotic species demonstrated different HRP preference patterns (Figs. S7A-D). PolyS containing sequences were commonly shared, while *Homo sapiens* exhibit a higher proportion of polyP and polyA in HPS-Pos proteins compared to others (Figs. S7A-D). Strikingly, the unicellular protozoan parasite *Plasmodium falciparum*, which causes malaria in humans, has an exceptionally high proportion of polyN-containing proteins as its only prominent HRP population (Fig. S7B, F), while their functions remain unclear. Furthermore, *Saccharomyces cerevisiae* exhibits a higher proportion of polyQ-containing proteins (Fig. S7A), indicating a robust protein quality control system that confer exceptional resilience to protein aggregation.

**Figure 3.**
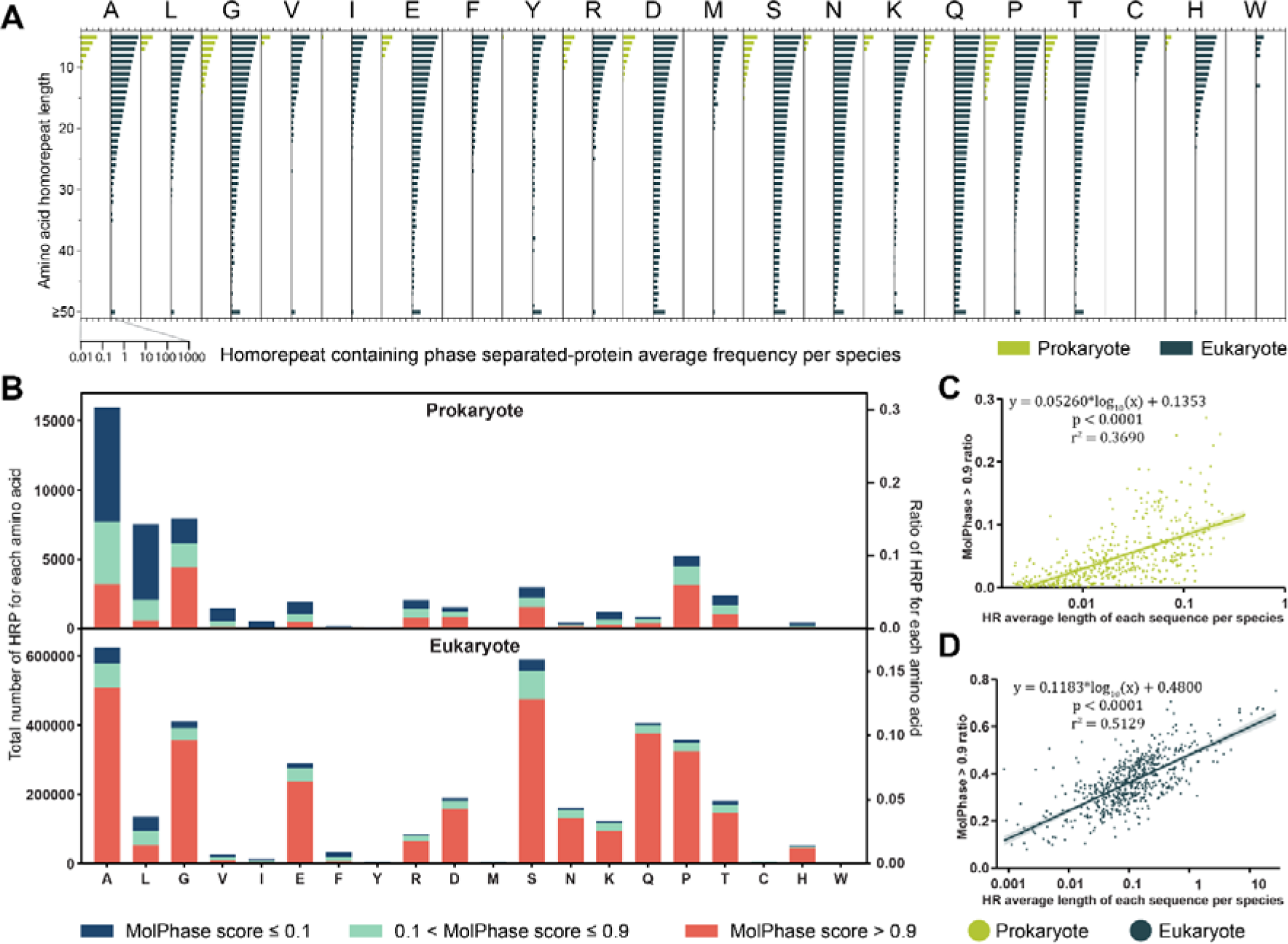
Homorepeat amino acid analysis in prokaryotes and eukaryotes proteome. (A) Distribution of the length of repeat in HRP. The x-axis indicates the average occurrence frequency per species of certain homorepeat length within the entire prokaryotic or eukaryotic domain, and the y-axis indicates the repeat length of consecutive amino acids. The minimum repeat length is 5 amino acids. The repeat equal to or longer than 50 amino acids were added together and plotted as the last bar. The detailed calculation methods are listed in the materials and methods section. (B) The distribution of HRP for each amino acid in prokaryotes and eukaryotes. The left y-axis represented the total amount of HRP for each amino acid, and the right y-axis represented the ratio of HRP for each amino acid. The HRP counts for each species, categorized as prokaryotic or eukaryotic, were summed together and plotted based on the amino acid type in the homorepeat. (C)-(D) Non-linear regression of HR average length of each protein sequence per species with HPS-Pos ratio in (C) prokaryotes and (D) eukaryotes. The demonstration of x-axis was under log10 transformation.

To further elucidate the relationship between homorepeat and PS, we next plotted the average homorepeat length against the HPS-Pos ratio. Both prokaryotic and eukaryotic species exhibit an overall positive correlation between the length of homorepeats and PS propensity (Figs. 3C-D). This suggests that increased PS propensity may provide assembly flexibility to counterbalance the potential toxicity associated with HRPs. While interestingly, compared to the linear regression results (r^2^ in prokaryotes = 0.3357, eukaryotes = 0.1772), the log_10_ transformation of HR average length in each sequence (r^2^ in prokaryotes = 0.3690, eukaryotes = 0.5129) showed better fitting (Figs. 3C-D), indicate the accumulation of homorepeat toxicity is slow. Protein abundance analysis of HRPs in four representative eukaryotic species suggested that proteins with long homorepeats tend to maintain a relative low abundance (Figs. S7E-H), especially those for aggregation-prone PLD amino acids (Q, N, Y), which also hinted the high expression level of HRPs might be toxic to cells and tightly controlled is needed. Interestingly, in *Saccharomyces cerevisiae*, both highly abundant short homorepeat-containing proteins and less abundant long homorepeat-containing proteins are predominantly stress-adaptation-responsive proteins, suggesting HRP-mediated condensation for stress adaptation instead of only introducing aggregation toxicity.

Amino acid homorepeats in proteins can be deleterious to cellular function and are frequently implicated in human diseases; for example, the abnormal expansion of glutamine repeat would lead to human Huntington’s disease (*28*). Organisms may need to evolve multilayered regulatory mechanisms to balance their requirements with toxicity risks. By analysing multiple factors including relative translational rate, protein abundance, protein half-life of HRP in *S. cerevisiae*, Chavali *et al.* found that HRP are more stringently regulated than non-HRP in synthesis, activity and degradation (*26*). Another research reported that highly disordered proteins are more stringently controlled than low disordered proteins (*29*). To further dissect the regulatory of PSPs, we analyzed the protein abundance in all HPS-Pos and HPS-Neg, which revealed that HPS-Pos proteins tend to maintain a lower level compared to HPS-Neg proteins (Figs. S8A-D). We further utilized the available data from budding yeast to analsyse PSPs. The HPS-Neg have a higher ratio of long poly(A) tail-containg proteins (Fig. S8E), which could enhance the transcriptional initiation (*30*). Whereas HPS-Pos proteins were translated less and slower (Figs. S8F, G). Furthermore, the HPS-Pos proteins have a higher turnover rate (Fig. S8H), suggesting faster protein degradation and consistent with their nature of rich in post-translational modifications (*31*). Combined with the abundance data of HRPs in four representative species (Figs. S7E-H), the production and maintenance of PSPs are tightly regulated across different kingdoms of life.

### PhaseHub: integrating quantitative proteomics, interactome, and biophysical features to identify multicomponent condensation hubs

Homotypic PS depends on intrinsic biophysical properties and surrounding environmental conditions, whereas cellular condensations often involve heterotypic interactions. Protein concentration and the capacity for intermolecular interactions among components of a heterotypic phase separation system are critical for determining the efficiency of scaffolding and the hierarchical organization of multicomponent condensates when they start to arise (*32*). Here, we integrated MolPhase predictions for homotypic PS with experimental data on protein abundance and interactions of four eukaryotic model species, *Saccharomyces cerevisiae*, *Plasmodium falciparum*, *Homo sapiens,* and *Arabidopsis thaliana,* to identify multicomponent condensation hubs in diverse biological processes (Figs. 1C-F). We retrieved the protein abundance information from PaxDB 5.0 (*33*) and interaction data from IntAct (*34*) and STRING (*35*). The 80^th^ percentile of interaction partner number and protein abundance were set as thresholds to classify the HPS-Pos protein. By crossing the number of protein interaction partners and protein abundance together, the abundant hub proteins were classified (Fig. 4), as these proteins are hot targets for serving as PS functional scaffold hubs. Protein family analysis revealed that RNA-binding domain, RNA recognition motif domain, and helicase showed high frequency in these abundant hub proteins (Figs. 5A-D, I). Combined with its filter condition of high abundance and interactors, it further emphasizes the prevalence of nucleic binding domain in promoting PS *in vivo*. Overlapping of KEGG pathway enrichment analysed of abundant hub proteins in four species demonstrated five pathways were significantly enriched, with further emphasize the important of PS in nucleic acid related pathways (Figs. 5E-H, J).

**Figure 4.**
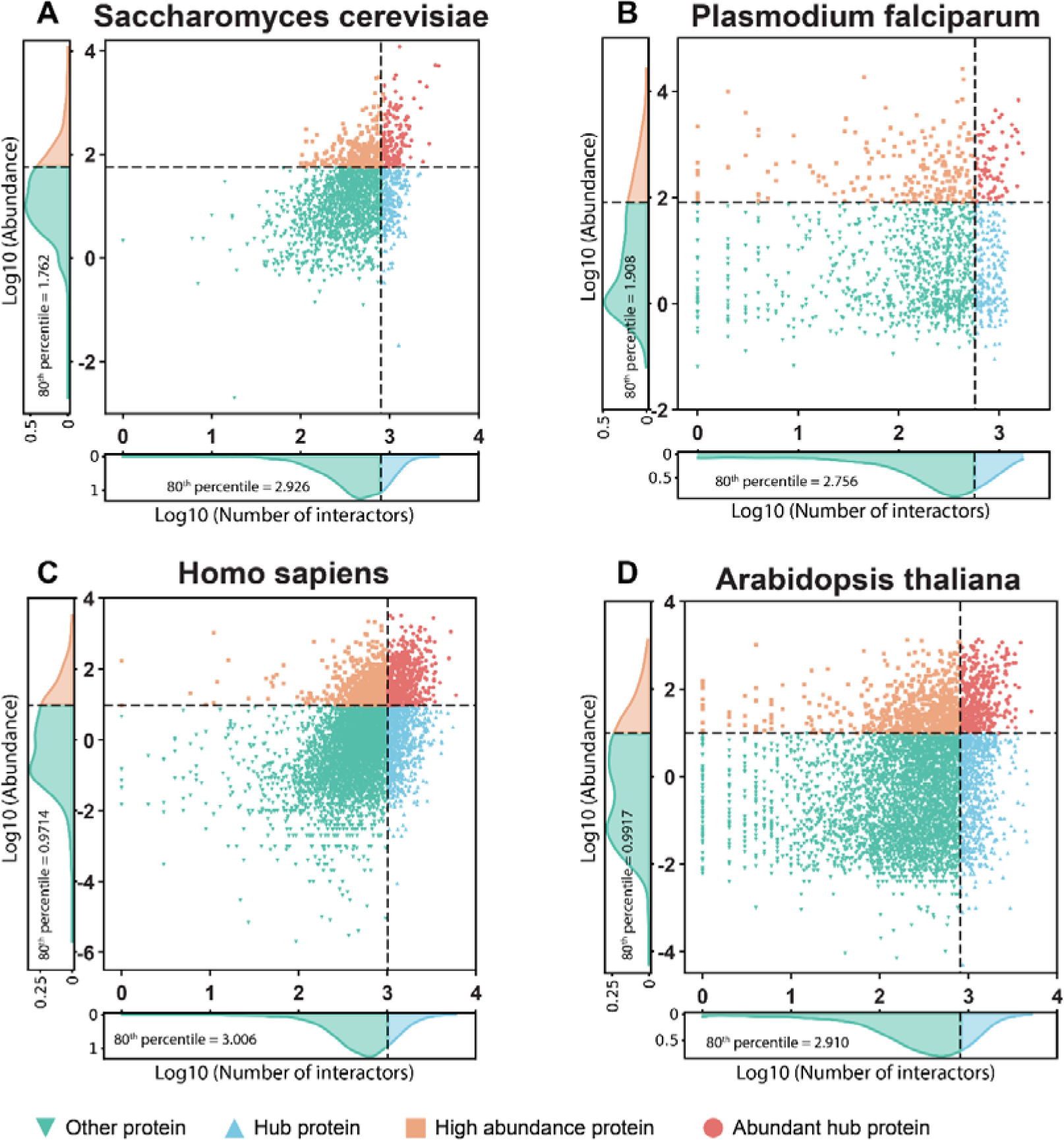
Identification of the abundant hub protein in HPS-Pos protein within four representative species. (A)-(D) Overlapping of protein abundance and protein interactome of HPS-Pos protein in (A) *Saccharomyces cerevisiae*, (B) *Plasmodium falciparum*, (C) *Homo sapiens*, and (D) *Arabidopsis thaliana*. Proteins of the top 20^th^ percentile of abundance were label as ‘high abundance protein’, and proteins of the top 20^th^ percentile of the amount of interactor label as ‘hub protein’. The distinct proteins were identified based on overlapping the top 20^th^ percentile of high abundance protein and hub protein, labeled as ‘abundant hub protein’.

**Figure 5.**
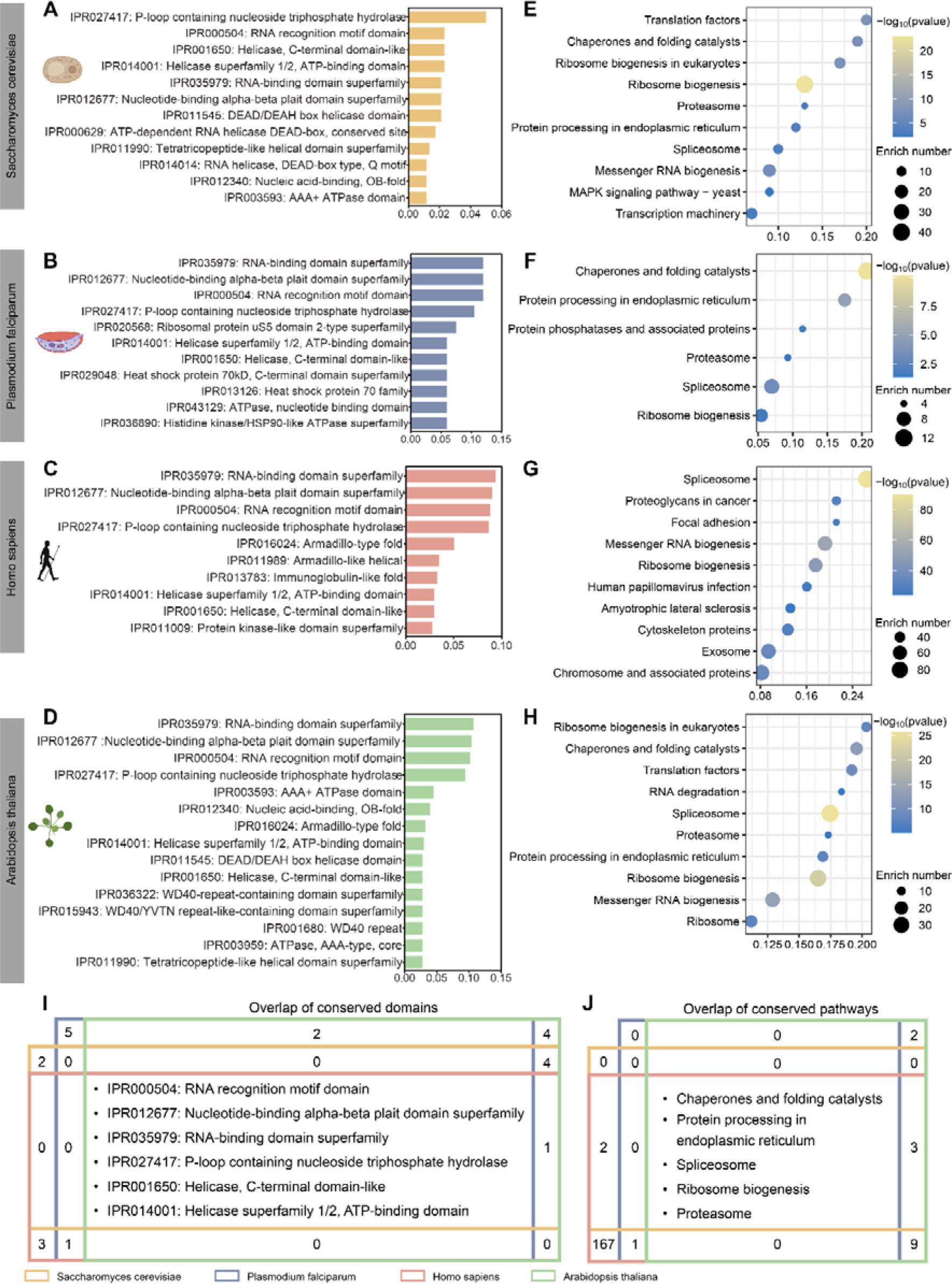
KEGG pathway and protein family enrichment of abundant hub phase-separated protein in four representative species. (A)-(D) Protein family frequency of abundant hub protein in (A) *Saccharomyces cerevisiae*, (B) *Plasmodium falciparum*, (C) *Homo sapiens*, and (D) *Arabidopsis thaliana*. Protein family information was retrieved from InterPro. (E)-(H) KEGG pathway enrichment analysis for HPS-Pos proteins with abundant hub protein in (E) *Saccharomyces cerevisiae*, (F) *Plasmodium falciparum*, (G) *Homo sapiens* and (H) *Arabidopsis thaliana*, where the x-axis shows the −log_10_ p-value for the Fisher exact test, the y-axis lists enriched pathways, the ratio indicates the percentage of the enriched item number in entire pathway, and the enrich number shows the number of enriched protein number in the pathway. For all the significant (p ≤ 0.05) enriched pathways, the top 10 were shown here. (I) Overlapping of top enriched protein family shown in Figure 5A-D. (J) Overlapping of all the significant (p ≤ 0.05) enriched pathways of four representative species.

To facilitate the visual exploration and analysis of multicomponent phase separation systems for discovering functional hubs for the proteins of interest across species, we next developed PhaseHub, an interactive computational platform. PhaseHub integrates quantitative proteomics, interactomics, and homotypic PS propensity and its biophysic feature analysis from four representative eukaryotic species. Using direct searches or BLAST with protein sequences (Figs. 6A, B), users can visualize potential co-condensation partners for each protein, customized by filtering PS-propensity (MolPhase score). Results are displayed based on cellular abundance and interaction capabilities, highlighting their potential roles as scaffolding or client components homorepeat (Figs. 6C, D). Additionally, each protein candidate is listed with a clickable ID, enabling easy access to its annotation and sequence, with directly highlighted homorepeat. These interactors of phase-separated scaffold proteins are hot candidates for clients recruitted into biomolecular condensates (*19*), the interactors list might provide a useful list for client identification (Figs. 6E, F). Targeting these hub interactors offers an alternative approach for engineering condensation properties and biochemical activities. Besides, the physiological protein concentration data could also help users to understand whether the protein abundance aligns with its functional roles and PS propensity inside the cells. The cross-talks among potential multicomponent condensates can also be directly observed, positioning them as the most promising factors, as additional scalfolders or functional clients. In summary, the PhaseHub serves as a data-driven platform for generating hypotheses of functional condensation based on protein interaction network mapping and identifying crucial pathway nodes.

**Figure 6.**
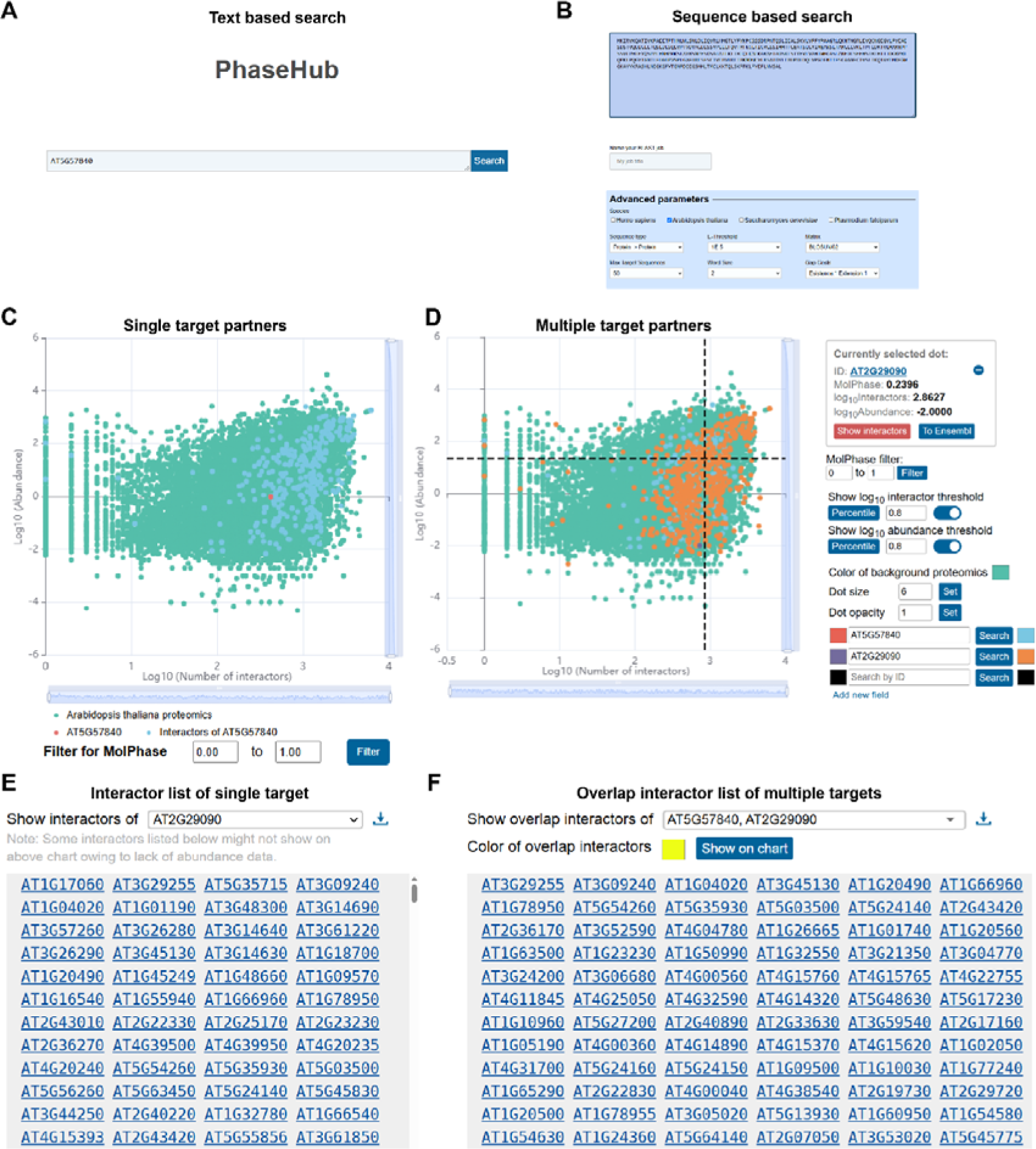
Overview of PhaseHub. (A)-(B) The searching function of PhaseHub, (A) text based searching and (B) sequence-based searching. (C)-(D) The interaction targets network demonstration for (C) single target and (D) multiple target partners. For (C), the interaction partners of *Arabidopsis thaliana* protein AT5G57840 were demonstrated and for (D), the interaction partners of AT5G57840 and AT2G290290 were demonstrated. (E)-(F) The interaction partners list for (E) single target and (F) multiple targets. For (E), the interaction partners of *Arabidopsis thaliana* protein AT5G57840 were listed and for (F), the overlapped interaction partners of AT5G57840 and AT2G290290 were listed.

## Discussion

PS could utilize diverse strategies to regulate biochemical reactions and versatile cellular activities, including concentration, specificity, and components to recruit or exclude functional modules (*36*). Our study systematically explored the biophysical features of phase separation across 1106 species, comparing them with protein abundance, interactome, and HPR in model species to elucidate potential multicomponent phase separation hubs (PhaseHub) and their functional evolution in living organisms.

### Phase separation directs the evolution of molecular grammar in life complexity

The conserved stability of protein folding is essential for preserving geometric information at protein interfaces, ensuring precise molecular interactions. However, this stability can restrict the structural plasticity and multivalency required for adaptive functionality, which are critical for development and resilience. Multivalent macromolecular condensation are often tunable to generate functional activities on depamand, by achieving dynamic interactions for signaling processes, moving beyond the traditional lock-and-key binding model (*37–39*). Thus, the functional evolution of phase separation is more than identity the amino acid mutations within structured domains, as emphasized in traditional evolutionary analyses. It also encompasses the preservation of biophysical property patterns that are essential for maintaining dynamic multivalent weak interactions, allowing continuous material exchange with the surrounding environment. PS can thus coordinate functional outcomes in orchestrating various processes using key catalytic modules. For example, the multivalent scaffolding protein LAT can form different signaling hubs and amplify the signal by interacting with various adapters during T-cell signaling (*40–45*). The flexibility and adjustability given by phase separation confer organism a useful tool to modulate the more and more complex life, which manifested as the increasing of phase separation with the gaining of genome size (Fig. 2).

In our study, we analyzed multicomponents phase separation by two key properties— protein abundance and associative capabilities—which do not always coincide, though we identified a substantial subset of proteins exhibiting each trait within the proteomes of model organisms. Abundance is primarily governed by evolutionary optimization of transcription, translation, and protein homeostasis, whereas associative features and valency arise from sequence-based evolutionary selection. By integrating insights from protein biophysical properties, molecular grammar and multivalent protein-protein interactions (PPIs), we constructed a systems-level analysis of PS. By examining over 1000 species across the evolutional tree and specific analysis of four eukaryotic model species with experimental data-derived PPI and abundance information, we identified the biological processes that heavily rely on PS to facilitate functional assemblies in each species and those conserved across different kingdoms of life (Fig. 5). Additionally, the highly interactive scaffolding proteins and their potential hub components in these pathways could be analyzed by PhaseHub. A user-friendly interface integrating protein abundance, interactome, and biophysical features offers an intuitive framework for hypothesis generation, enabling researchers to investigate multicomponent, multivalent condensation hubs in signaling pathways. This platform would accelerate hypothesis development with comprensive feature analysis, facilitating rational design for studying functional macromolecular condensation. Because these properties may represent distinct strategies with synergy for leveraging phase separation in biological functions. For instance, high abundance could gate specific signaling events to form responsive condensates during development and stress responses, while varying associative motifs with differing affinities might regulate condensation via hierarchical assembly. With the evolution from simple to complex, organisms evolve network-like biomolecular systems to cope with environmental changes and developmental demands, accompanied by a dramatic increase in the number of proteins involved in more cellular processes. The expansion of protein-coding genes coupled with the rise in PSP ratio enhanced the capabilities to form adaptive molecular assemblies, which could respond to diverse surrounding enviroments on demand. For example, liquid-like stress granules could form inside the cells upon stress conditions such as heat shock, which were assembled by stalled mRNA, translation initiation factors and other proteins, and could act as plugs at endomembrane damage sites and enable rapid repair (*46*). At the same time, the homotypic PSPs could also serve as sensors for environmental stimuli. For example, the *Arabidopsis* SEUSS protein could form liquid-like condensation under osmotic stress and promote stress tolerance gene expression (*47*). In summary, stress-triggered PS is an adaptive response to cope with environmental clues (*48*), and the bomb of PSPs from prokaryote to eukaryote is reasonable for evolutional adaptation (Fig. 1).

### Phase separation is a double-edged sword for life

Dynamic condensation of functional hubs could aid in organizing the more complex life activities by spatiotemporally regulated on demand. However, the plateau in the increase of PSP in eukaryote suggests that the potential detrimental effects may counterbalance their functional benefits. Especially the phase transition from soluble liquid-like status to amyloid, which is the cause of various diseases (*49, 50*). Since the formation of PS is conditional dependent, the vulnerable droplet state is sensitive to intrinsic and extrinsic condition changes and undergoes a phase transition to aggregation (*51*). The maturation from fluidic droplets to the amyloid state will lead to protein dysfunction and toxicity, which has been proven to be associated with various diseases (*52*). For some proteins such as FUS, even single amino acid mutation could greatly accelerate aggregation formation and is associated with neurodegeneration (*50*). The PSPs mutation lead to dysfunction would further exacerbate as cells aging, insoluble protein content is likely to accumulate due to the deterioration of physiochemical conditions and cell aging (*53*). And the progressively accumulation of aggregation upon aging is accompanied by a gradual loss of the compartmentalization conferred by PS (*54*). Worse still, the homorepeat can intensify the aggregation property (*55*), which are enriched in PSPs (Fig. 3B), further emphasize the necessity of restrict PSP prevalence in life.

The potential detrimental effects of PSPs may be the primary reason preventing simple prokaryotes from acquiring more PSPs (Figs. 1A-J). Conversely, this suggests that eukaryotic systems have evolved effective safeguard mechanisms to mitigate aggregation toxicity, such as the tight regulation at the transcriptional, translational, and proteostasis levels (Fig. S8). Considering the energetic constraints may explain why prokaryotes cannot sustain complex systems akin to those of eukaryotes. As protein turnover is an energy-intensive process in living organisms, consuming over 38% of the total energy produced by the lactic acid bacterium *Lactococcus lactis* and roughly 20% of the human resting metabolic rate (*56, 57*). Due to the absence of endosymbiotic mitochondria, prokaryotic cells rely solely on the plasma membrane for chemiosmotic ATP synthesis, converting proton motive force into ATP. However, this process becomes significantly less efficient as cell volume increases, as the surface area-to-volume ratio decreases, limiting energy availability (*58*). In contrast, eukaryotic cells overcome this constraint through the presence of mitochondria, which provide a highly efficient and scalable energy production system via oxidative phosphorylation across their internal membranes (*58*). The energy surplus generated by mitochondria enables eukaryotic cells to synthesize PSPs without the energetic constraints faced by prokaryotes. This may explain why eukaryotes, even with genome sizes comparable to those of prokaryotes, exhibit a higher ratio of HPS-Pos (Figs. 2D, H). Consequently, the fundamental differences in energy production mechanisms—plasma membrane-based in prokaryotes versus mitochondria-driven in eukaryotes—underpin the significant disparity in PSP abundance between them.

In evolutionary biology, the “C-value paradox” refers to the absence of a direct relationship between an organism’s biological complexity and its genomic DNA content (*59*). This paradox likely reflects a balance between genome size and the availability of sufficient multimodular gene functions. Intriguingly, we found that in prokaryotes and eukaryotes with smaller genomes—where genome size remains below a critical threshold needed for diverse gene utilization—the expansion of genes related to phase separation positively correlates with biological complexity, which does not fall under the “C-value paradox”. However, as genome size increases over evolutionary time, the flexibility and diversity of PSPs also rise. Beyond a certain point, if sufficient modularity exists to adapt to various conditions, further increases in genome size may become unnecessary, reflecting a trade-off between energy expenditure and functional efficiency. Yet, diverse life cycles and environmental pressures can impose constraints on genome and PSP expansion, potentially introducing detrimental effects. Understanding the regulation dynamics of “C-value paradox in phase separation” requires future studies.

## Material and Methods

### Proteome information collection and cleaning

All the proteome sequence data sources are listed in Table S1. Since the minimum sequence length limitation for MolPhase is 40, and the extremely long amino acid sequences will consume too many computational sources. Therefore, all the sequences longer than 10000 amino acids or shorter than 40 amino acids will be removed. Sequences that contained illegal characters out of 20 basic amino acids, such as “X” will also be filtered. In eukaryotic cells, the gene transcription process is often affected by different conditions, and other transcripts are formed through alternative splicing. Thereby, one gene could produce multiple protein products. We used the gene transfer format files, which were retrieved with proteome files together to map the alternative splicing product for each gene, and only the longest protein isoform for one gene would be kept for further analysis. The cleaned proteome would be used for MolPhase prediction and all the others analysis in this study, including PS promoting feature analysis, amino acid homorepeat analysis, and KEGG/protein family enrichment analysis, etc.

### Phase separation prediction and biophysical, biochemical features evaluation

Protein phase separation prediction was carried out by MolPhase (*12*). The protein biophysical and biochemical features used in this study, including IDR percentage, pi interaction, PLD likelihood, and LCR percentage, were predicted by respective tools, as mentioned before (*12*).

### Regression analysis

For genome size information used in the regression analysis, the genome size was acquired from NCBI genome assembly statistics. Owing to the assembly data missing, only partical species were used for genome size correlation analysis. In total, 132 archaea, 364 bacteria, 156 fungi, 118 protists, 174 animals and 122 plants were used. Regression analysis was performed using the least squares method in GraphPad Prism 9.5.0.

### Amino acid homorepeat analysis

In this article, the amino acid homorepeat was recognized as at least five or longer identical consecutive amino acids in the sequence. The cutoff was based on previous research that (i) the occurrence of five or longer continuous stretches was statistically significant compared to randomly generated sequences (*26*), and (ii) the known shortest homorepeat with length variation associated with human diseases is a five residues polyaspartate (polyD) stretch in cartilage oligomeric matrix protein, which leads to pseudoachondroplasia and multiple epiphyseal dysplasias (*60*). The identification of homorepeats and counting of amino acid frequency were carried out by Python script based on the proteome files. The homorepeat length in HRP distribution frequency for each amino acid in Fig. 3A was calculated as follows:

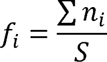

where *f_i_* represents the normalized frequency of protein with i-length consecutive repeats per species in respective prokaryotic or eukaryoteic species, *n_i_* is the number of proteins containing i-length consecutive repeats in a single species’ proteome, which would sum from all prokaryotic or eukaryotic species, and S is the total number of species in prokaryotic or eukaryotic groups.

And the homorepeat average length of each sequence per species in Figs. 3C and D was calculated as follows:

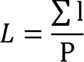

where L represents the average length of homorepeat in one species, l represents the length of homorepeat in one sequence, with sum of all homorepeat of 20 amino acids in one species, and P represents the total number of protein sequences in one species.

### Protein abundance and protein interactome analysis

Protein abundance was retrieved from PaxDB 5.0 (*33*), the integrated abundance data of whole organisms were retrieved and used as the representative abundance for each protein. Protein binary interaction data was collected from IntAct (*34*) and STRING (*35*). The binary interaction pairs were combined from IntAct and STRING, follow by removing repeat pairs. Proteins with invalid ID or interacted with exogenous protein such as pathogentic proteins were removed. Then the interaction partners were counted for each protein. Only the HPS-Pos protein with both available abundance and interaction partner number will be used for enrichment analysis.

### Protein enrichment analysis

For Kyoto Encyclopedia of Genes and Genomes (KEGG) enrichment analysis, KEGG background files of each species were obtained from KEGG GenomeNet (*61*). Protein of interest were mapped respective KEGG pathway k-number by eggNOG (*62*). Protein family information was classified by InterPro (*63*). Significant test was performed by Fisher’s exact using the Python package, SciPy (*64*). Bubble plots of KEGG enrichment analysis were plotted by SRplot (*65*).

## Author contributions

Conceptualization: Q.L. and Y.M. Methodology: Q.L. and Y.M. Data curation: Q.L. Formal analysis: Q.L. and Y.M. Visualization: Q.L. Investigation: Q.L. and Y.M. Resources: Q.L. and Y.M. Writing—original draft: Q.L., W.G. and Y.M. Supervision: W.G. and Y.M. Funding acquisition: W.G. and Y.M.

## Data and materials availability

All data needed to evaluate the conclusions in the paper are present in the paper and/or the Supplementary Materials. The MolPhase prediction result of the proteome of 1106 species was available at https://phasehub.sbs.ntu.edu.sg/download.php.

## Supplementary Materials

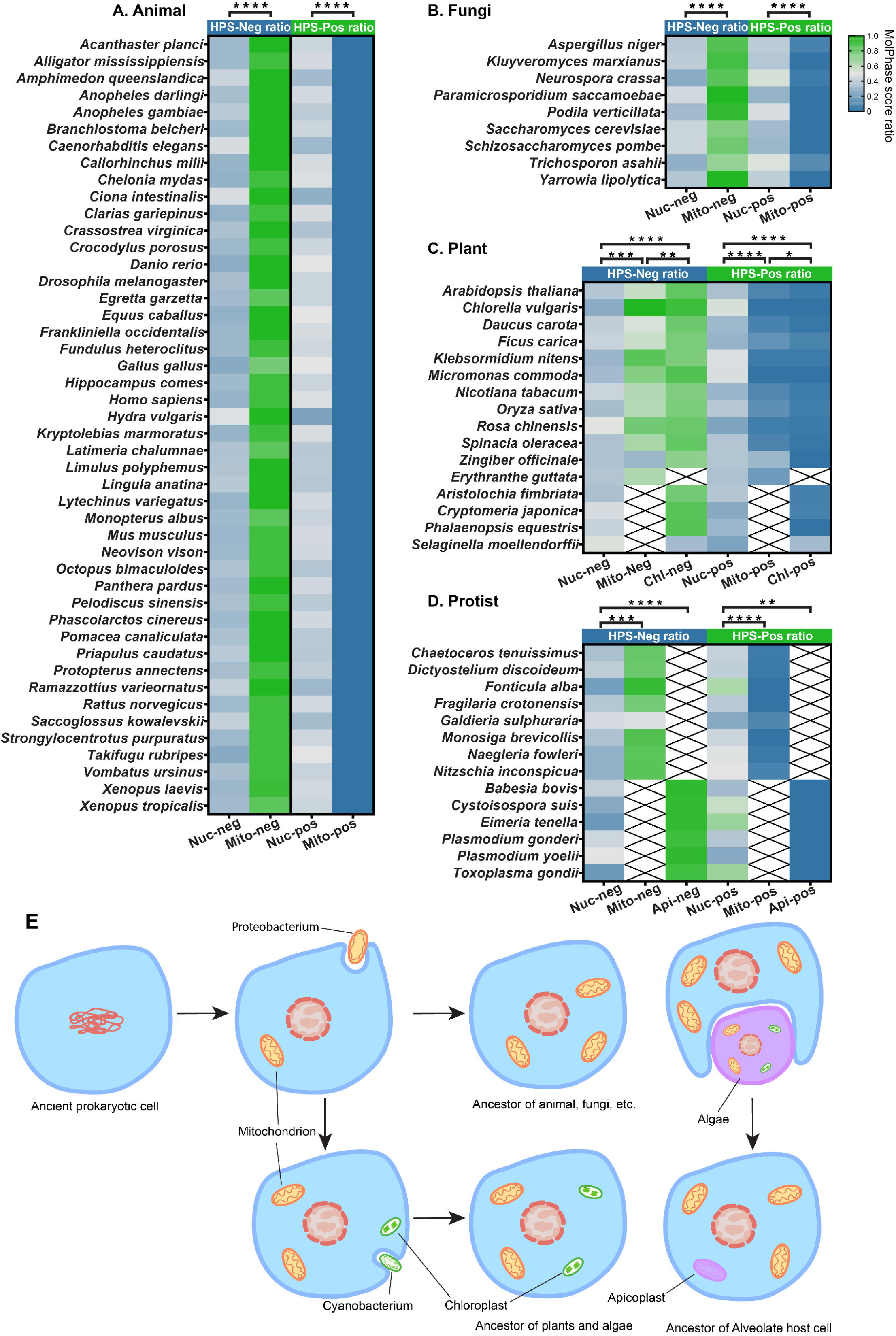

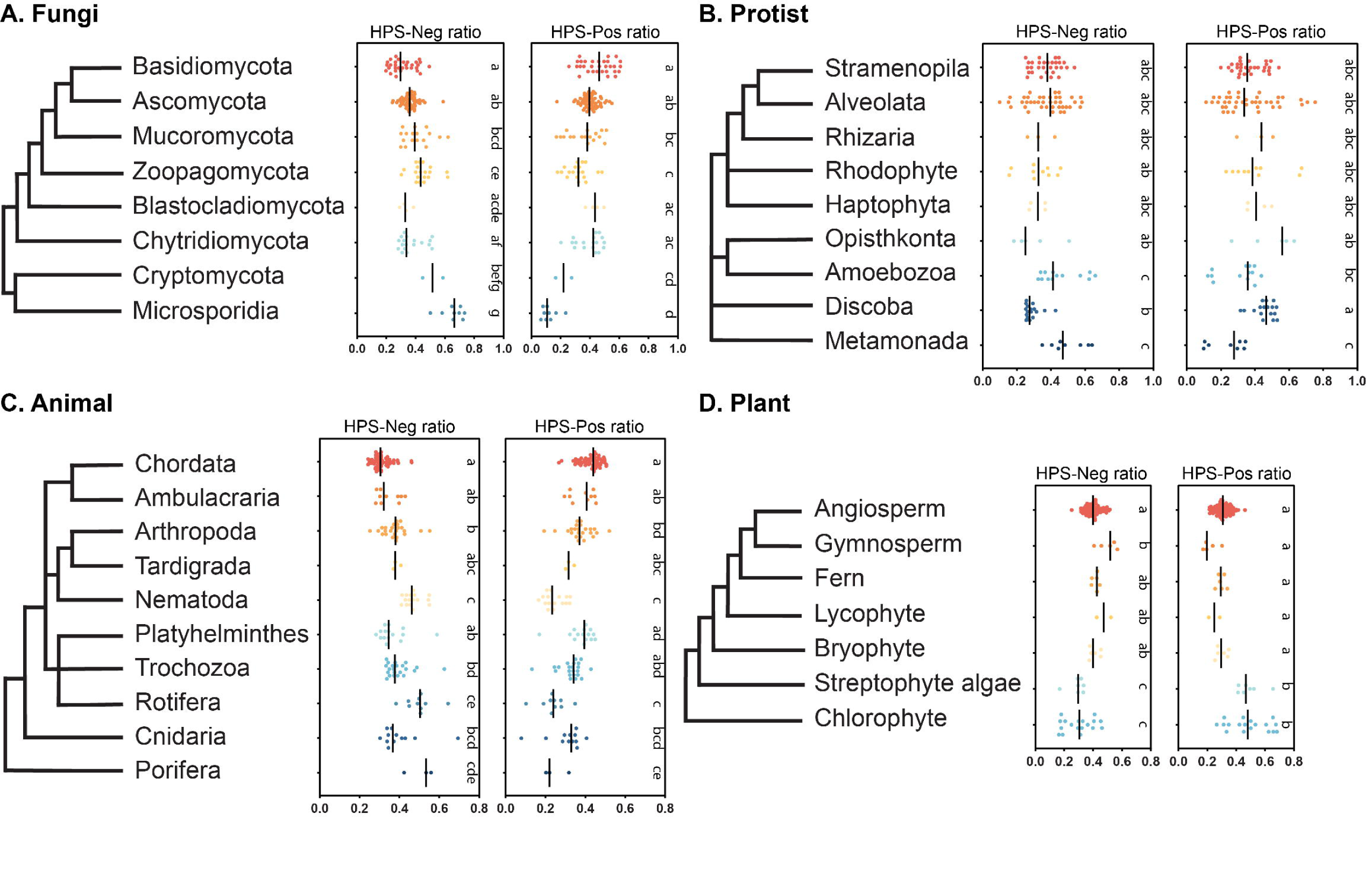

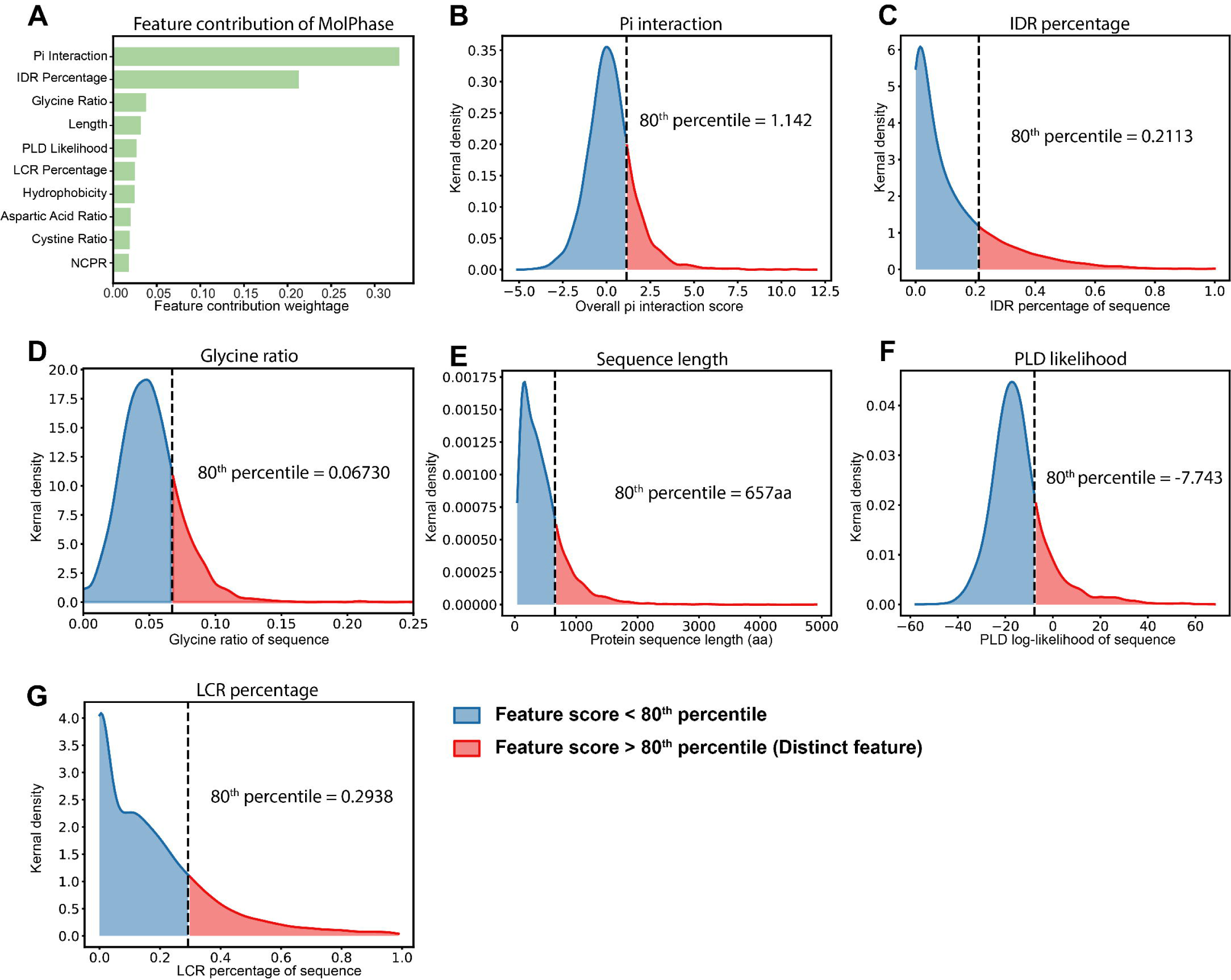

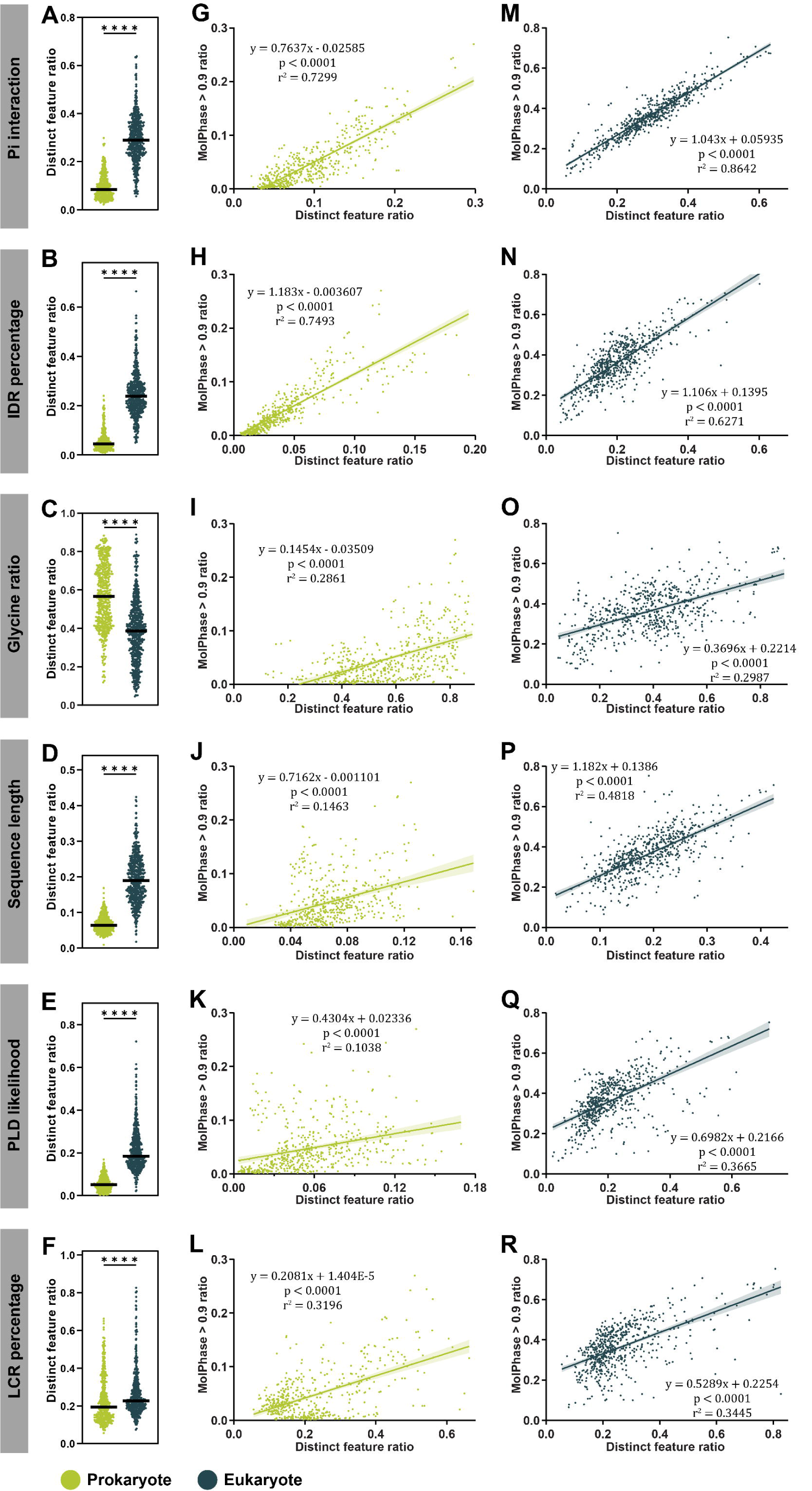

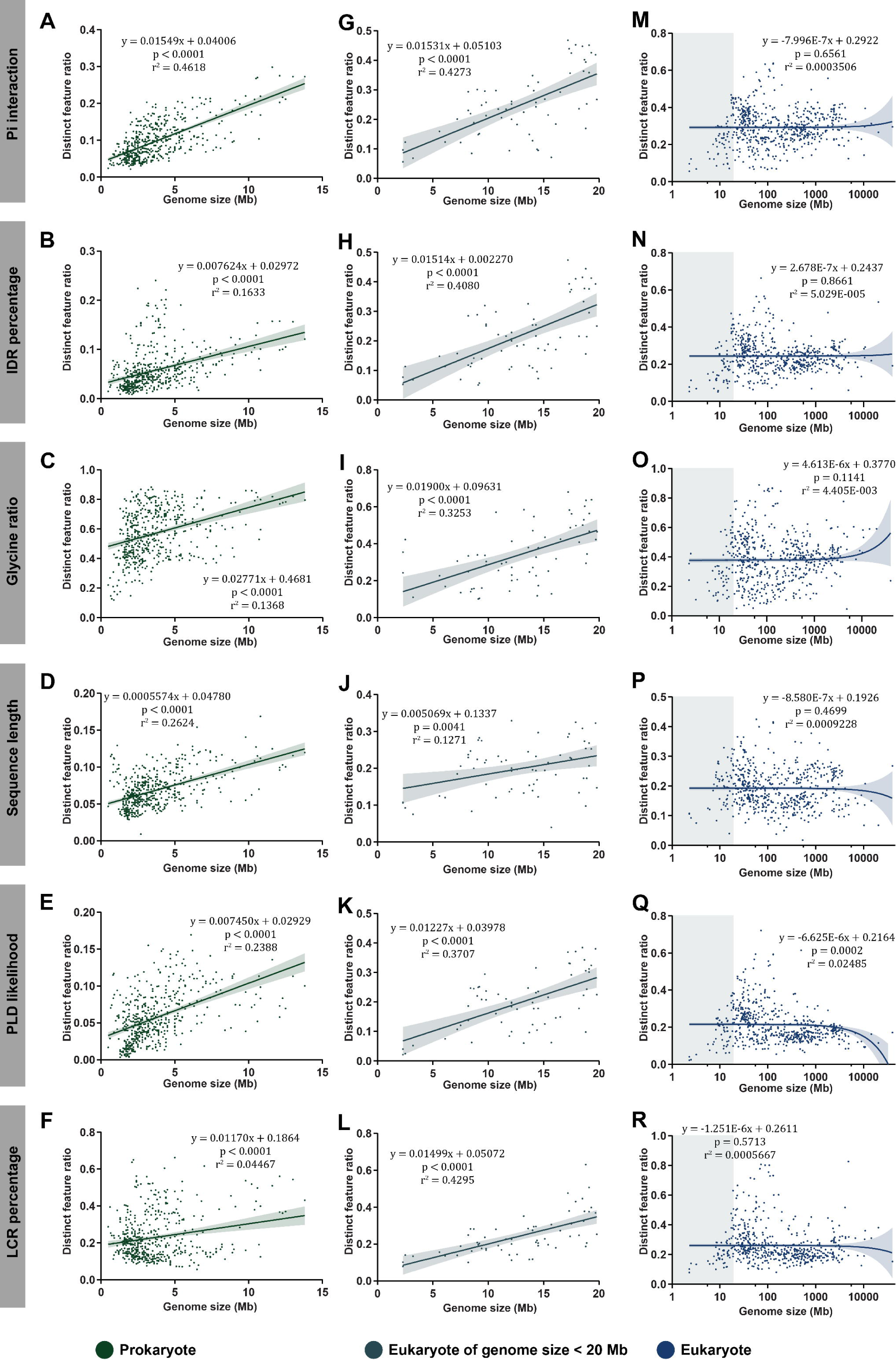

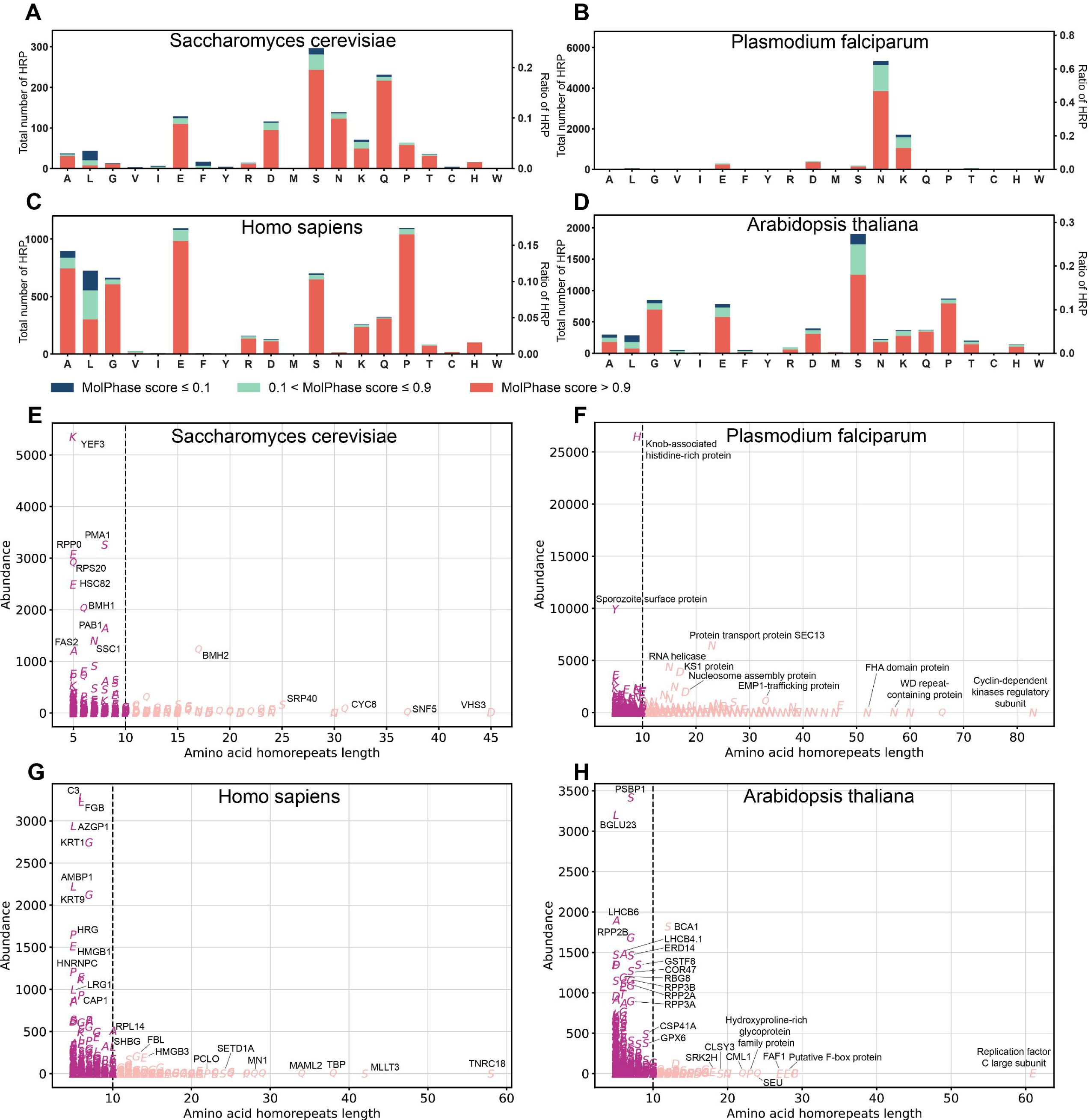

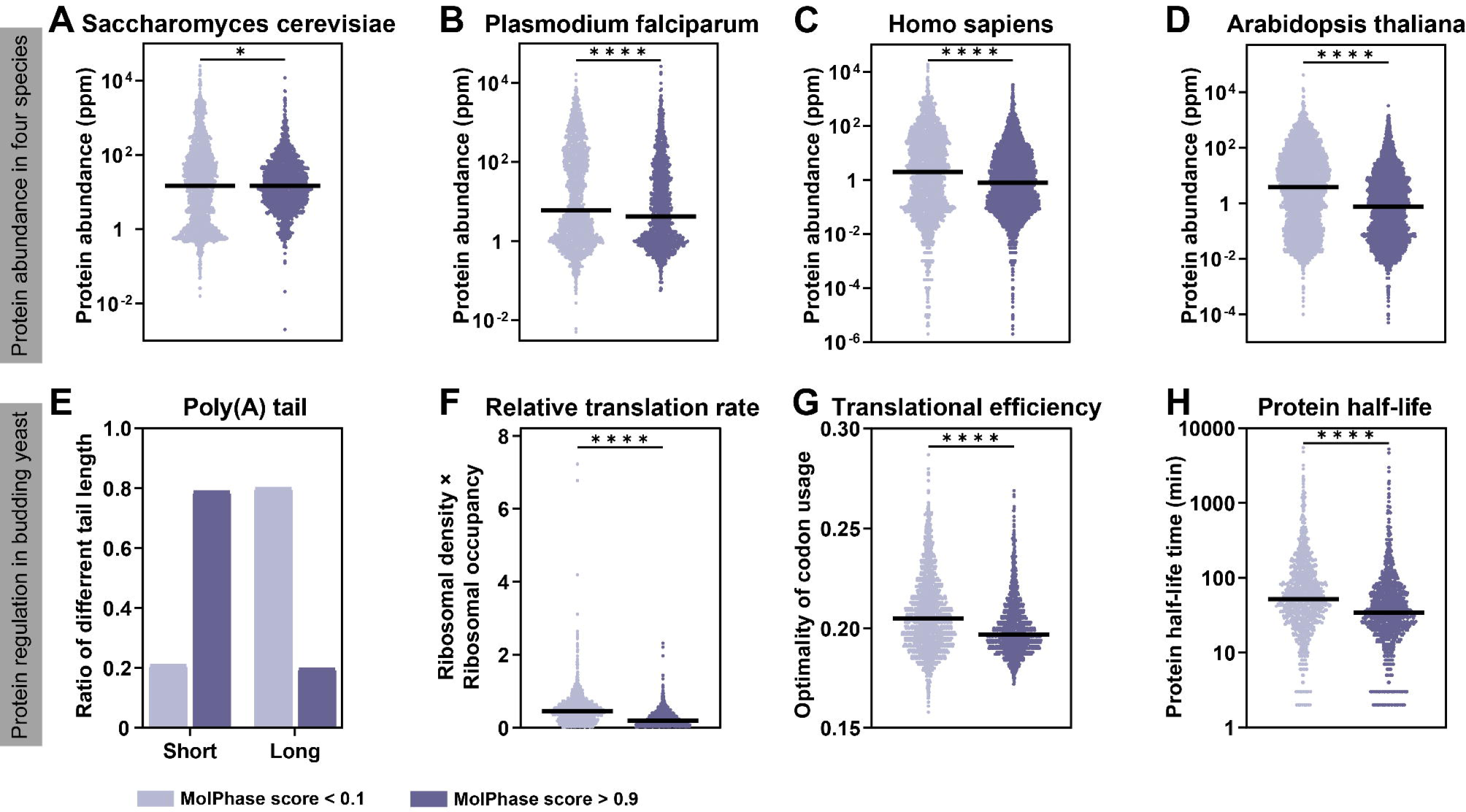

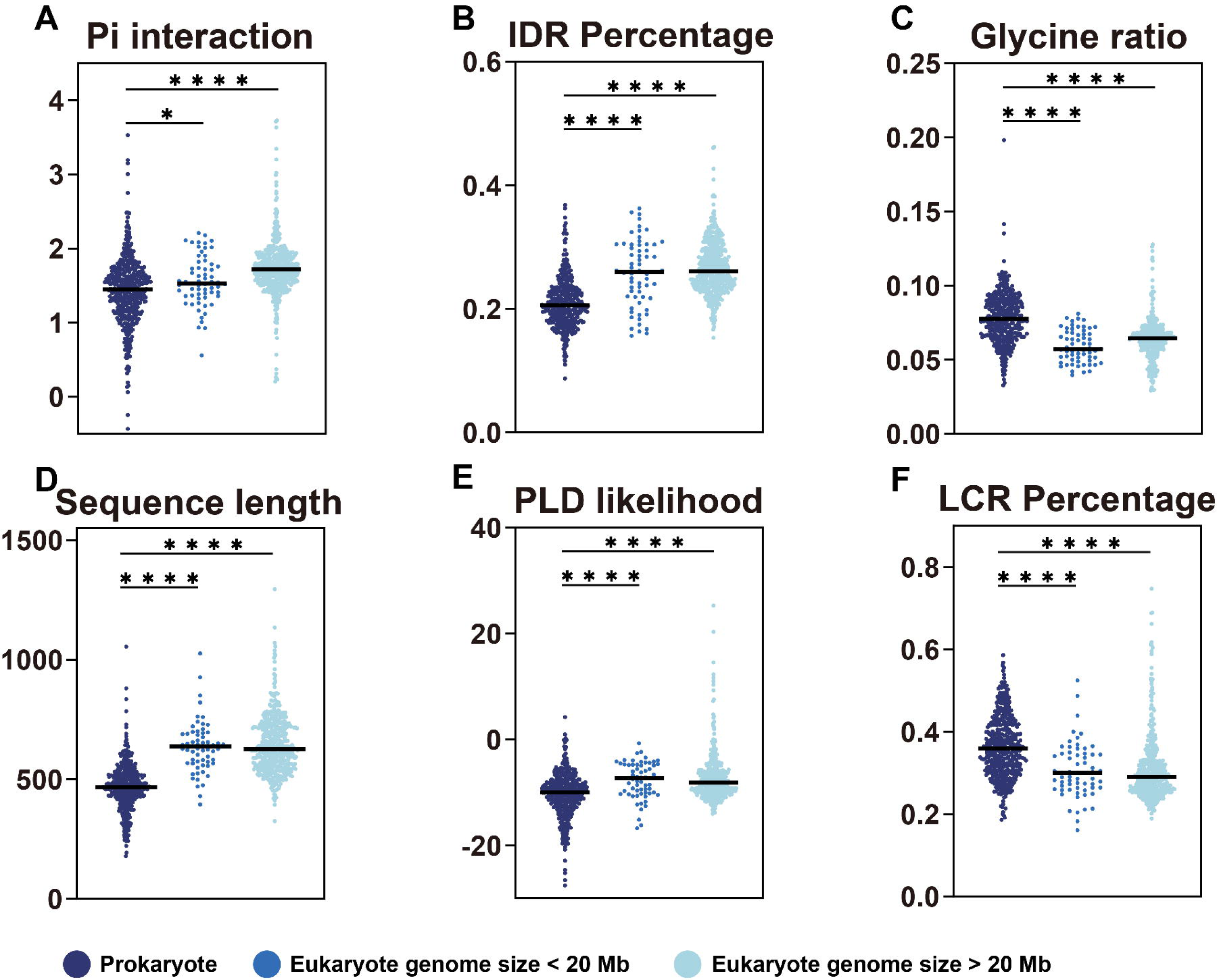

Table S1

